# Approximating Signal Sources in Stereo-EEG Single Pulse Electrical Stimulation using Re-referencing and Spectral Analysis

**DOI:** 10.1101/2025.03.03.641282

**Authors:** Michael A. Jensen, Harvey Huang, Nicholas M Gregg, Klaus-Robert Müller, Dora Hermes, Kai J. Miller

**Author notes:** Department of Neurosurgery, 200 1st Ave, Mayo Clinic, Rochester, MN, USA.

## Abstract

Stereoelectroencephalography (sEEG) is a method of epilepsy monitoring that uses stereotactically placed depth electrodes. Connectivity between brain regions may be assessed by delivering a current pulse from sEEG electrode(s) and monitoring all other electrodes for Brain Stimulation-Evoked Potentials (BSEPs). However, the shape, magnitude, and spectral content of BSEPs are sensitive to the analysis approaches used to study them. In order to understand the nuance between analysis and interpretation of BSEP data, we carefully apply and contrast modified common average & bipolar re-referencing techniques, combined with wavelet-based spectral analysis on measured data from 4 human subjects. Comparison between analysis approaches allows us to identify recording sites within, near, or far from the tissue source of the BSEPs they record. We further explore the interaction between re-referencing techniques and dipole source orientation, providing methods to estimate source location and upper bound of far field recordings. This systematic application and comparison of different analysis tools can enable those who study BSEPs to localize potential sources within, near, or far from measurement sites, and more accurately constrain how brain connectivity is understood from stimulation experiments.

**Author summary:** Brain stimulation evoked potentials can be driven by sources at, near, or far from the recording site. Classification of these evoked potentials into one of these domains can be done using established re-referencing and spectral analysis techniques. Specifically, modified common average and bipolar re-referencing allow delineation of near and far field boundaries while spectral analysis allows for identification of channels within active tissue.

## Introduction

Intracranial single pulse electrical stimulation (SPES) allows for the exploration of white matter connections between brain areas by stimulating one brain area and measuring evoked potential changes (BSEPs – brain stimulation evoked potentials) everywhere else. In addition to determining network maps, BSEPs have also been used more subtly - to interpret the directed nature of cortical connections [1], as well as their speed [2] and strength [3]. Each of these studies has made the assumption that the brain phenomena they are exploring are tied directly to the site of measurement. However, it has never been established how local these responses actually are, since BSEPs may represent conduction from more distant sources.

SPES is performed using either electrocorticography (ECoG) surface grids [1, 2, 4] which stimulate and record the tissue underneath, or stereoelectroencephalography (sEEG) depth electrodes [5–8]. ECoG has a natural relationship perpendicular to the cortical surface, while sEEG stimulate and record from tissue in all directions [9]. While both ECoG and sEEG record from surrounding and farther sources, sEEG’s sparse, volumetric nature increases the volume of potential sources without a compensatory increase in the density of recording sites. For example, a sEEG channel in the frontal operculum may record sources from insular, temporal, or ventral sensorimotor areas, which are all separated by sulci from one another. In this example, if stimulation of the motor thalamus were to generate a source in the ventral motor cortex, but the nearest recorded BSEP were in the operculum, a false conclusion could be drawn that the motor thalamus and operculum are connected.

The purpose of this work is to provide a framework and toolkit to assess where BSEP sources might originate in relation to the sEEG recording site. We do this by leveraging a set of clear re-referencing and spectral analysis techniques that are illustrated heuristically and also with patient sEEG data. Armed with this toolkit, the reader will be able to more clearly interpret their own BSEP data and constrain their hypotheses within distinct measurement domains.

## Methods and Materials

### Ethics Statement

The study was conducted according to the Institutional Review Board of the Mayo Clinic (IRB 15-006530), which also authorizes sharing of de-identified data. Each patient or parental guardian provided informed consent as approved by the IRB. All T1 MRI sequences were de-faced prior to publication using an established technique [10] to avoid potential identification (*Data and Code Availability*).

### Subjects

Four subjects (2 female, 878 total electrodes, 271 stimulation pairs) participated in our study, each of whom underwent placement of 13-18 sEEG electrode leads for seizure network characterization in the treatment of drug-resistant epilepsy (Table 1). Electrode implant locations were planned by the clinical epilepsy team based on typical semiology, scalp EEG studies, and brain imaging. No plans were modified to accommodate research, nor were extra electrodes added. All experiments were performed in the epilepsy monitoring unit or pediatric intensive care unit at the Mayo Clinic in Rochester, MN.

**Table 1.**
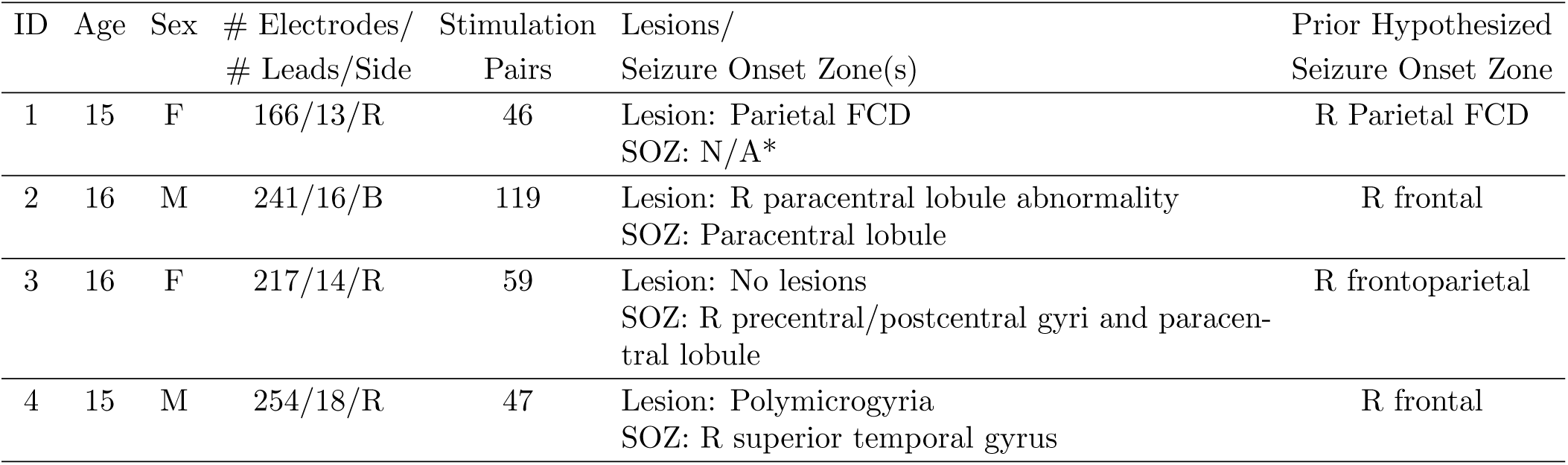
Subject Information: Age, Sex, Number of Contacts/Leads/Laterality, number of stimulation pairs, seizure onset zones and foci location, hypothesis prior to sEEG, seizure onset zone hypothesized prior to sEEG implantation. *indicates no stereotypical seizures were recorded during the EMU stay.

| ID | Age | Sex | # Electrodes/<br># Leads/Side | Stimulation<br>Pairs | Lesions/<br>Seizure Onset Zone(s) | Prior Hypothesized<br>Seizure Onset Zone |
| --- | --- | --- | --- | --- | --- | --- |
| 1 | 15 | F | 166/13/R | 46 | Lesion: Parietal FCD<br>SOZ: N/A* | R Parietal FCD |
| 2 | 16 | M | 241/16/B | 119 | Lesion: R paracentral lobule abnormality<br>SOZ: Paracentral lobule | R frontal |
| 3 | 16 | F | 217/14/R | 59 | Lesion: No lesions<br>SOZ: R precentral/postcentral gyri and paracentral lobule | R frontoparietal |
| 4 | 15 | M | 254/18/R | 47 | Lesion: Polymicrogyria<br>SOZ: R superior temporal gyrus | R frontal |

### Single Pulse Electrical Stimulation (SPES) and sEEG Recordings

Subjects were originally stimulated as part of two other studies such that stimulation sites were chosen based on pre-selected hypotheses. Each electrode pair was stimulated 12 times, with a single 6 mA, 100µs, biphasic pulse, and a random 3-7 s interval between pulses, using a g.Estim Pro electrical stimulator (gTec, Schiedlberg, Austria). Voltage data were recorded on a g.HighAmp amplifier at 4800 Hz. An electrode located in white matter was selected as the hardware ground. Electrodes within the clinically-determined seizure onset zones were excluded from analysis. Individual response trials containing artifacts or epileptiform activity were identified and excluded. An epileptologist was present during all stimulation sessions to monitor ictal activity, with seizure rescue drugs prepared in case stimulation led to seizures. Data were processed using custom MATLAB code (see *Data and Code availability*). Signals were high-pass filtered above 0.5 Hz in forward and reverse directions with a second-order Butterworth filter and notch filtered at 60, 120, and 180 Hz (*±*1Hz width).

### Common Average Re-referencing by Least Anticorrelation (mCAR)

Data were first re-referenced using a modified common average method. This technique minimizes the anticorrelation between any single channel and the remaining re-referenced signals. Beginning with the least covariant channel, channels are iteratively added to create a reference signal that is compared to the remaining channels left out. The subset averaged signal with minimal correlation, by simple correlation, to the remaining channels not included in the subset averaged signal is selected for re-referencing. This method, termed Common Average Re-referencing by Least Anticorrelation (CARLA) [11], is the specific instsance of a modified common average re-referencing (mCAR) used throughout the manuscript.

### Bipolar Re-referencing

Data were separately re-referenced in a bipolar fashion by taking the difference between voltage time series of adjacent channels. This produced channels that reflect mixed activity between the potential measured at two adjacent electrode recording sites. Plotted points of both stimulation sites and bipolar re-referenced recording sites represent an interpolated point between the two electrodes that make up each pair. Only immediately adjacent channels were used to generate pairs - i.e. 3.5mm center-to-center from one another on the same lead.

### Electrode Localization and Anatomic Labeling

For each subject, the pre-operative T1 MRI was realigned to the anterior and posterior commissure stereotactic space (AC-PC) [12], and then co-registered to the post-implant CT using SPM12 [13]. Each electrode was localized in AC-PC space using the CT scan.

Channels were plotted on axial, coronal, and sagittal planes of the T1 MRI using the SEEGVIEWpackage [6] to show evoked response locations in an interpretable, clinically familiar manner. It should be noted that as each electrode is projected to the nearest slice from its position in AC-PC space, a site shown in gray matter may artifactually appear to be in white matter after projection, and vice versa.

T1 MRIs for each subject were segmented using the built in autosegmentation algorithm from Freesurfer 7 [14]. Labels were assigned to each subject’s gyral and sulcal anatomy during autosegmentation using the Destrieux cortical atlas [14]. Regions containing the string “Thalamus” were labeled as thalamic and regions containing the string “White” were labeled as white matter (Supplementary Figs 1,2).

### Canonical Response Parameterization (CRP)

The CRP method was applied to recorded evoked potentials from 0.015 s to 1 s after stimulation [15]. Briefly, the CRP method is a machine learning technique that examines each stimulation-recording site interaction to identify a reproducible shape, and if it exists, quantifies its duration. This response shape can be regressed out of singular trial responses to obtain a residual signal (Fig 1, Supplementary Fig 1). In this study, wavelet-based spectral analyses were applied to this residual signal component because stimulation-induced asynchronous activity, reflecting broadband power spectral changes, will differ between trials and therefore sit within the residual signal. These asynchronous, broadband changes are associated with changes in neural firing rate [16].

**Fig 1.**
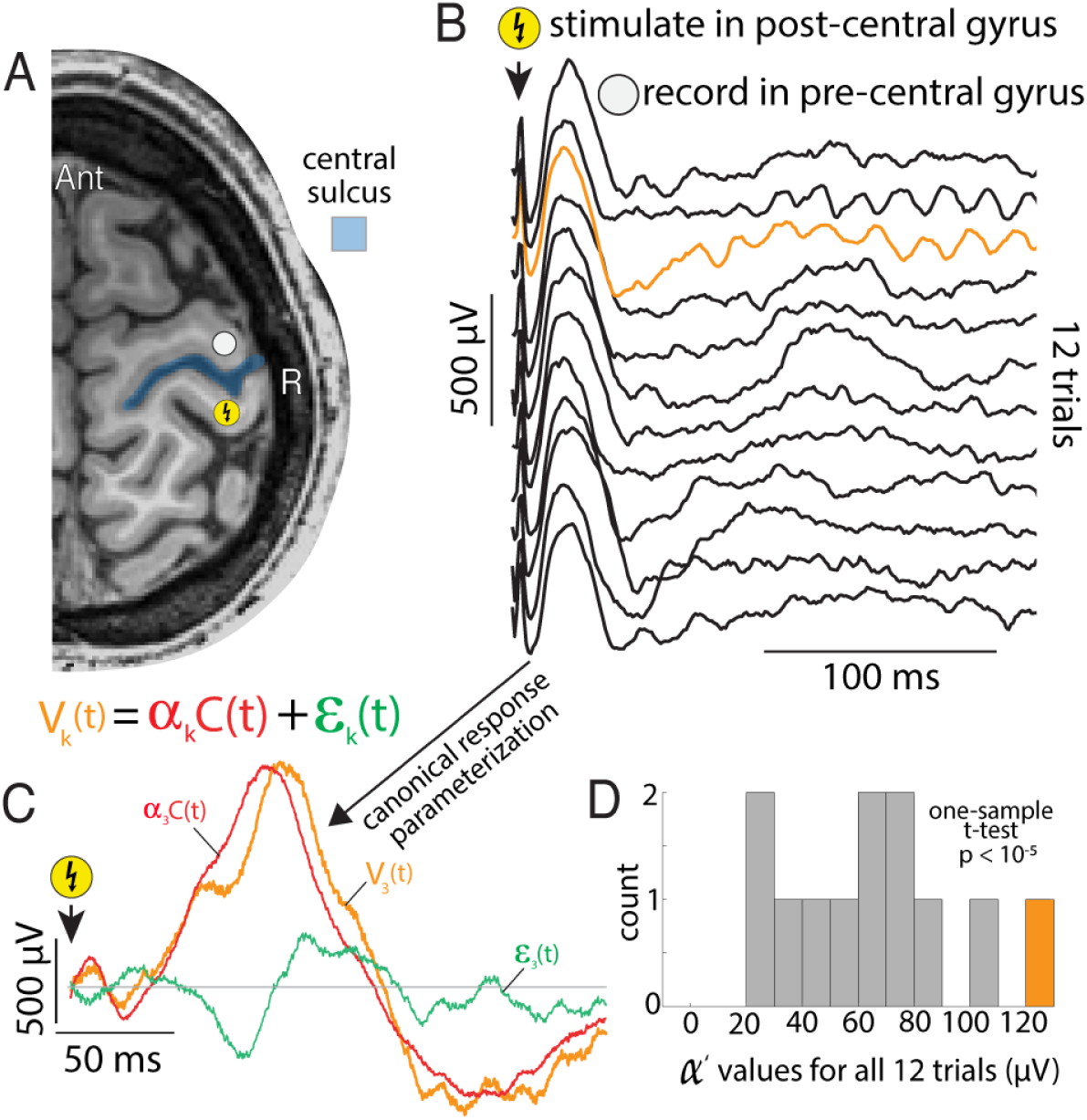
Single pulse electrical stimulation (SPES) to assess connectivity between brain regions, (data from Subject 1). **A.** An example SPES was delivered to the post-central gyrus while local field potentials were recorded from the pre-central gyrus. **B.** Stimulation of the post-central gyrus induces voltage deflections in the pre-central gyrus of a stereotyped form (often called a cortico-cortico evoked potential – CCEP). **C.** SPES induced voltage deflections across all stimulation trials can be parametrized using the canonical response parametrization (CRP) method. This CRP method takes as input all SPES induced voltage deflections and identifies the duration of significance in response (*τ_R_*) in addition to a canonical response shape (C(t) – red line) across all trials. Each trial is then compared to C(t) to identify the magnitude of the response (*α*) and residual noise (green line). **D.** The significance of an stimulation-recording site interaction is quantified using a one-sample t-test, comparing the distribution of normalized *α^′^* values to zero.

For each stimulation-recording site interaction, the significance of the evoked response is assessed using a one-sample t-test between the recorded *α*’ values across all stimulation trials and zero. p-values less than 0.05 were considered significant (Fig 1).

### Generation of synthetic evoked voltage deflections, broadband power increases, and background noise

*Evoked Raw Voltage Deflections:* synthetic evoked voltage deflections of 200 ms in duration were synthesized by taking the product of a Gaussian, a 1 Hz since wave, and 1.5 Hz sine wave. 20 voltage deflections were spaced 3 seconds apart to simulate BSEPs during 0.33 Hz stimulation in the absence of background noise. The amplitude of each BSEP was jittered between 98 and 102 percent of the simulated BSEP to allow for natural variations in the response amplitude. Five of these evoked potential time series were generated to represent 5 synchronously firing units (with slight jitter added between them).

*Evoked Broadband Power:* : broadband power was modeled to represent the summation of asynchronously arriving 50 ms post-synaptic potentials from 6000 pre-synaptic neurons onto a single postsynaptic neuron (i.e. in the dendrites of a cortical pyramidal neuron). Each post-synpatic potential was created by taking the product 50 ms exponential decay and power functions, and was designed to arrive randomly in time with a randomly determined magnitude between -1 and 1 *µ*V [17]. A 2 Hz firing rate was added coincident with each BSEP (see *Data and Code Availability*). Five stimulation evoked broadband time series were generated, representing five stimulation-responsive asynchronously firing units.

*Background Physiological Noise:* asynchronous noise was generated using the same model as evoked broadband, but firing rate was held constant across all time points at 2 Hz. A single time series was generated to represent the cumulative background physiological noise seen with intracranial recordings.

*Background Instrument Noise:* instrument noise was generated by generating white noise with magnitude between -3 and 3 *µ*V.

### Stimulation Artifact Removal

Prior to the wavelet spectrogram calculations, each stimulation artifact was removed from each recorded time series to minimize the spread of artifact into the time-frequency bins surrounding stimulation. This was done by multiplying an inverted Hann window function with a base extended by 15 ms (”bathtub” window), suppressing the signal until 20 ms after stimulation onset (Supplementary Fig 1). Because early voltage deflections can be sharp, they often lead to undesirable artifact in spectra (Supplementary Fig 2). Thus, this window was chosen as we wish to simultaneously 1) minimize the likelihood of artifact, and 2) identify local cortical activity lasting longer than 20 ms.

### Calculation of Wavelet Spectrograms

Wavelet -based spectrograms were calculated by applying a 5-cycle Morlet wavelet [18] from 1 to 200 Hz to the post-stimulation voltage time series after the canonical response shape was regressed out of each trial (using CRP) leaving only the residual component, *ɛ*(*t*). When replacing the raw voltage, V(t), with *ɛ*(*t*), discontinuities may be introduced due to gaps between V(t) and *ɛ*(*t*) at the time points where *ɛ*(*t*) begins and ends (Supplemental Figure 2). To avoid discontinuities at the beginning of *ɛ*(*t*) if *C*(*t*) does not exactly go to zero, parameterization using the CRP method began at stimulation onset after stimulation rejection, assuring that V(t) and *ɛ*(*t*) start at the same value and are thus continuous. To avoid discontinuities at the end of *ɛ*(*t*), the upper bound of *τ_R_* was used. This upper bound is defined as the larger value between: 1) the time sample to the right of the original *τ_R_* that is 99% of the maximum cross projection magnitude at the original *τ_R_*, and 2) 10% of the difference between the projection magnitudes at *τ_R_* and the final time point 500 ms after stimulation. This provides a more generous estimate of *τ_R_*. For each time-frequency bin, power was quantified as the square of the amplitude of the wavelet transform of *ɛ*(*t*). Power at each Hz was normalized to the mean power from -220 ms to -20 ms prior to stimulation across all trials. See the supplemental material for a more in depth exploration of how best to quantify stimulation evoked spectra.

### Source Localization Using *α^′^* Values

In order to illustrate a back of the envelope approach to localizing and characterizing of the source of an evoked response, we use a case study in subject 3, with evoked responses calculated by mCAR. Evoked responses from 19 recording sites (4 - gray matter, 15 – white matter) in the mid-cingulate were re-referenced using a modified common average [11] and parameterized using a single canonical shape [15], (Fig 1), generating *α^′^* values for each site. Significant ERPs were determined by comparing the distribution of *α^′^* values of each channel to zero (Fig 1). As *α^′^* carries units of µV, the mean *α^′^* for each recording site served to estimate the electric potential around each site. Significant channels were considered to be within the influence of a common dipole.

Because the mean *α^′^* values across all stimulation trials and positions were known for each of the significant channels, the position and orientation of the source dipole could be approximated. Assuming a dipole source such that the *α^′^* values follow an inverse squared law (Coulombs’ law), the 3-D position of the source were estimated by minimizing the squared error between the *recorded* electric potential (*α^′^* values) and *modeled* electric potential at each channel using gradient descent, minimizing the objective function:

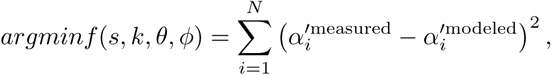

sequentially varying *s, k, θ, ϕ*, where

- *f* (*s, k, θ, ϕ*) is the objective function that depends on the estimated source position **s** and the proportionality constant *k* (treated as constant across space for this estimation),
- *N* is the number of channels,
- 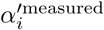 is the *α^′^* measured (*µV*) at the *i*th channel · normalized to the maximum *α^′^* across all channels,
- 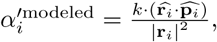
- 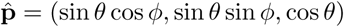 is the unit vector representing the dipole’s direction, with angles *θ* and *ϕ* representing the polar and azimuthal angles, respectively,
- 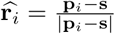 is the unit vector from the source **s** to the *i*th channel,
- |**r***_i_|* is the distance between the source and the *i*th channel,
- **p***_i_* = (*x_i_, y_i_, z_i_*) is the position of the *i*th channel,
- *s* = (*x_s_, y_s_, z_s_*) is the inferred position of the dipole source.

Note that a more complete model would not keep k as a constant, but would vary by tissue type using a finite element approach.

### Data and Code Availability

All data and code are available at Open Science Framework (DOI: https://osf.io/qsyug/) and GitHub (https://github.com/michaeljensen42/Approximating_source_distance).

## Results

Bipolar single pulse electrical stimulation (SPES) was delivered using stereoelectroencephalography (sEEG) depth electrodes while voltage time series were recorded in all non-stimulating channels. For each non-stimulating channel, voltage times series from 0.015 s to 1 s after stimulation were parameterized and significance was assessed using outputs of the CRP method (Fig 1 - Methods) [15]. Significant voltage deflections are referred to as brain-stimulation evoked potentials (BSEPs).

### Three domains of BSEP measurements

Using signal processing and re-referencing techniques, we can approximate three spatial relationships of BSEPs to stimulation-induced electric fields (Fig 2). They may be loosely interpreted as within the active tissue source, near field, or far field. *Within active tissue:* As broadband power (BB) directly correlates to local cortical firing [17, 19], BSEPs whose residual terms contain stimulation-locked BB changes sit within active cortex (BB+, BSEP_BP_ +, BSEP_CAR_ +). BSEPs without concomitant BB changes may measure near or far from the tissue source. To disentangle these, we can use the scale of the electrode spacing to define this transition. Sources significantly further will be canceled by by bipolar re-rereferencing but may be seen in mCAR re-referenced data. *Near field:* The distance at which the electric field changes at a smaller scale than the distance between adjacent recording sites represents near field recordings. In these cases, recorded potentials are non-zero (BSEP_CAR_ +) and the differences in recorded potentials in adjacent sites are significantly different from noise (BSEP_BP_ +). *Far field:* The distance at which the electric field changes at a greater scale than the distance between adjacent recording sites represents far field recordings in which the recorded potentials are non-zero (BSEP_CAR_ +) but the differences between adjacent sites become indistinguishable from noise (BSEP_BP_ -).

**Fig 2.**
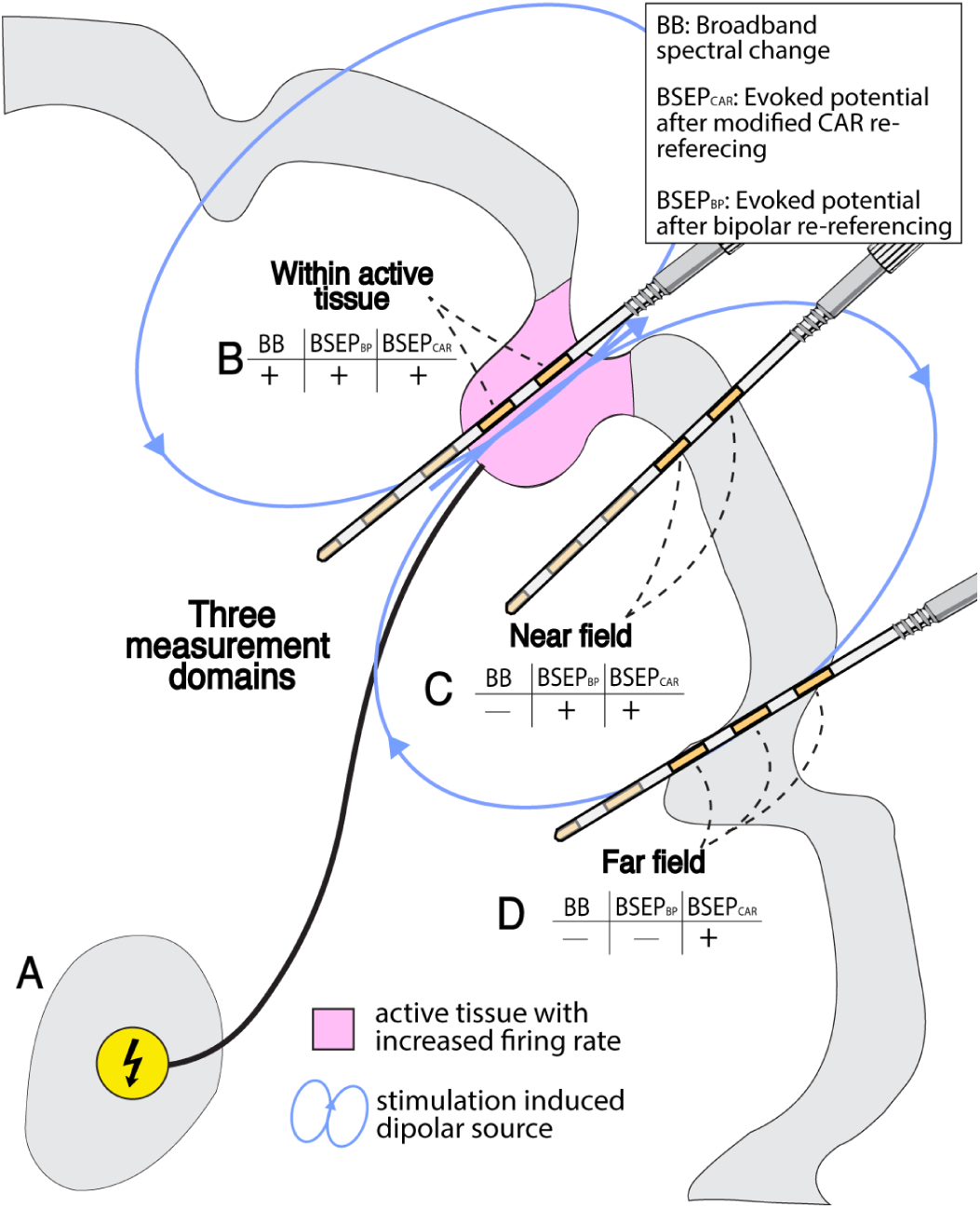
Three domains for measuring evoked responses. **A.** Stimulation of one site in the brain may induce propagation of a volley of action potentials to distant gray matter where many neurons are depolarized together, generating a dipole source(s). **B.** *Measuring within active cortex:* Recording sites may sit within the circuitry (gray matter) that is directly depolarized. This will result in recorded evoked potentials (BSEPs) whether using bipolar or a modified common average (mCAR) re-referencing technique. If stimulation induces activity of local neuronal circuitry, stimulation-evoked broadband power changes (BB) will be recorded as well. **C.** *Measuring near field:* Recording sites may sit near the source, but outside of the activated tissue. Due to the proximity of the recording sites to the source, local differences in electric field will result in measured BSEPs when using both mCAR and bipolar re-referencing. **D.** *Measuring far field:* Some recording sites may sit close enough to a source to record stimulation induced changes in field potentials, but far enough from the source such that spatial variation in potential is larger than the distance between adjacent contacts. Because of this, BSEPs will be seen using mCAR but not bipolar re-referencing. BSEP_BP_ = bipolar re-referenced BSEP, BSEP_CAR_ = BSEP re-referenced using a modified common average

These domains are illustrated in an example of stimulation in the right thalamus in subject 2, producing BSEPs in the central sulcus and adjacent white matter (Fig 3a). BB increases in addition to BSEPs can be seen in time-frequency spectrograms (Fig 3b-d) of the recordings from the central sulcus, suggesting this tissue is the cortical source of the recorded field. Two channels from the same lead sitting in white matter illustrate near and far domain responses revealed by the differences between mCAR and bipolar BSEPs (Fig 3e-j). Assuming that BB and BSEPs are due to asynchronous and synchronous processes respectively, we can generate synthetic data. These data show why BSEPs might be resolvable over greater distances than BB (Fig 4).

**Fig 3.**
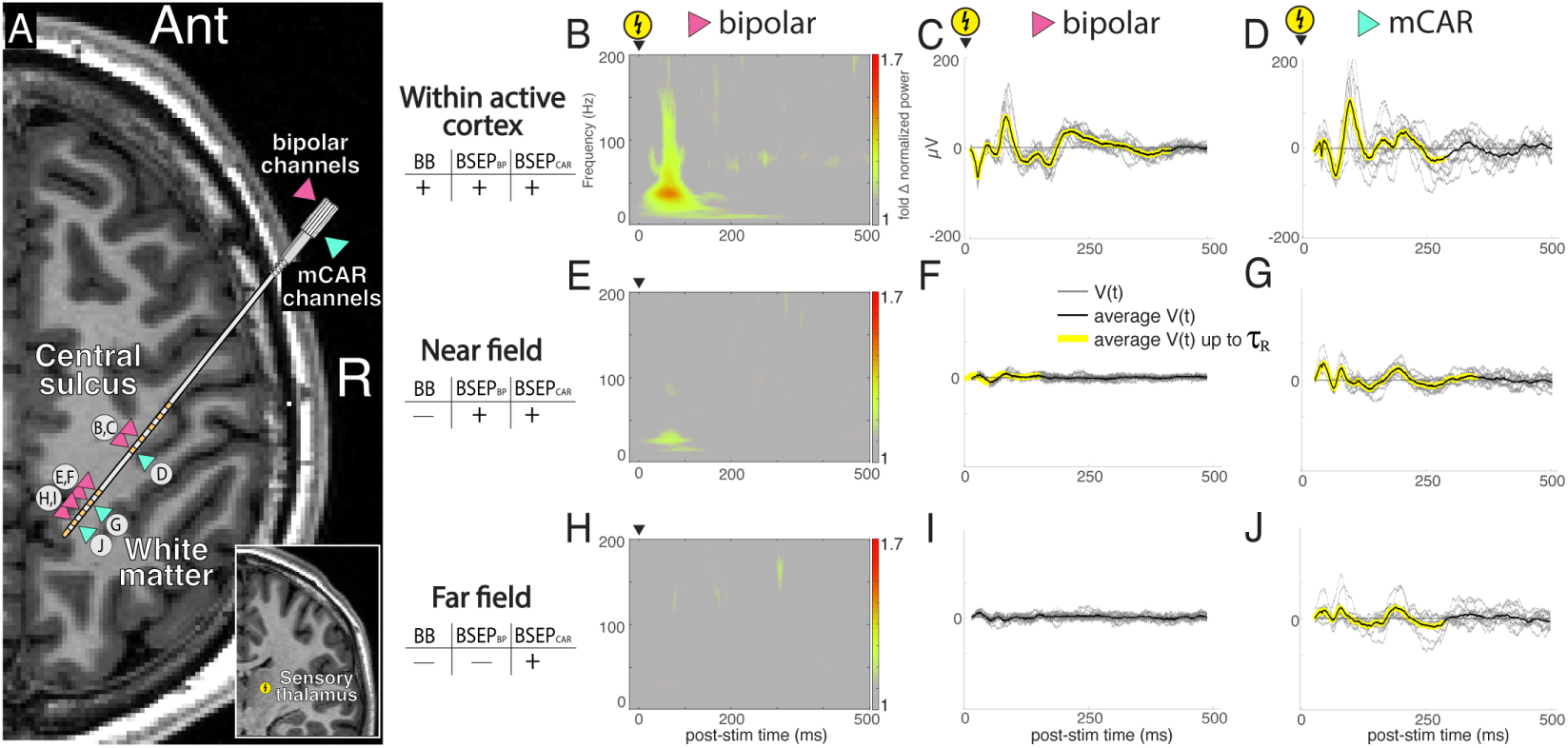
Example of the three domains of evoked responses, (data from Subject 2). **A.** Stimulation within the thalamus (coronal T1 MR inset - bottom right) leads to evoked responses in a lead recording running parallel to the central sulcus (axial T1 MR). **B-D.** *Example of measuring within active cortex:* significant mCAR and bipolar re-referenced evoked responses are recorded in the central sulcus contain asynchronous activity as seen by post stimulation increases in broadband power (most evident *>* 50 Hz) indicating that they are likely within active cortex. Note the white bars from 20 ms before until 20 after stimulation such that spectral artifact due to stimulation are not confused with true response. **E-G.** *Example of measuring near field:* significant mCAR and bipolar re-referenced evoked responses are recorded in white matter and lack asynchronous activity as seen by post stimulation broadband power. **H-J.** *Example of measuring far field:* significant mCAR but not bipolar re-referenced evoked responses are recorded in the white matter and lack asynchronous activity as seen by post stimulation broadband power.

**Fig 4.**
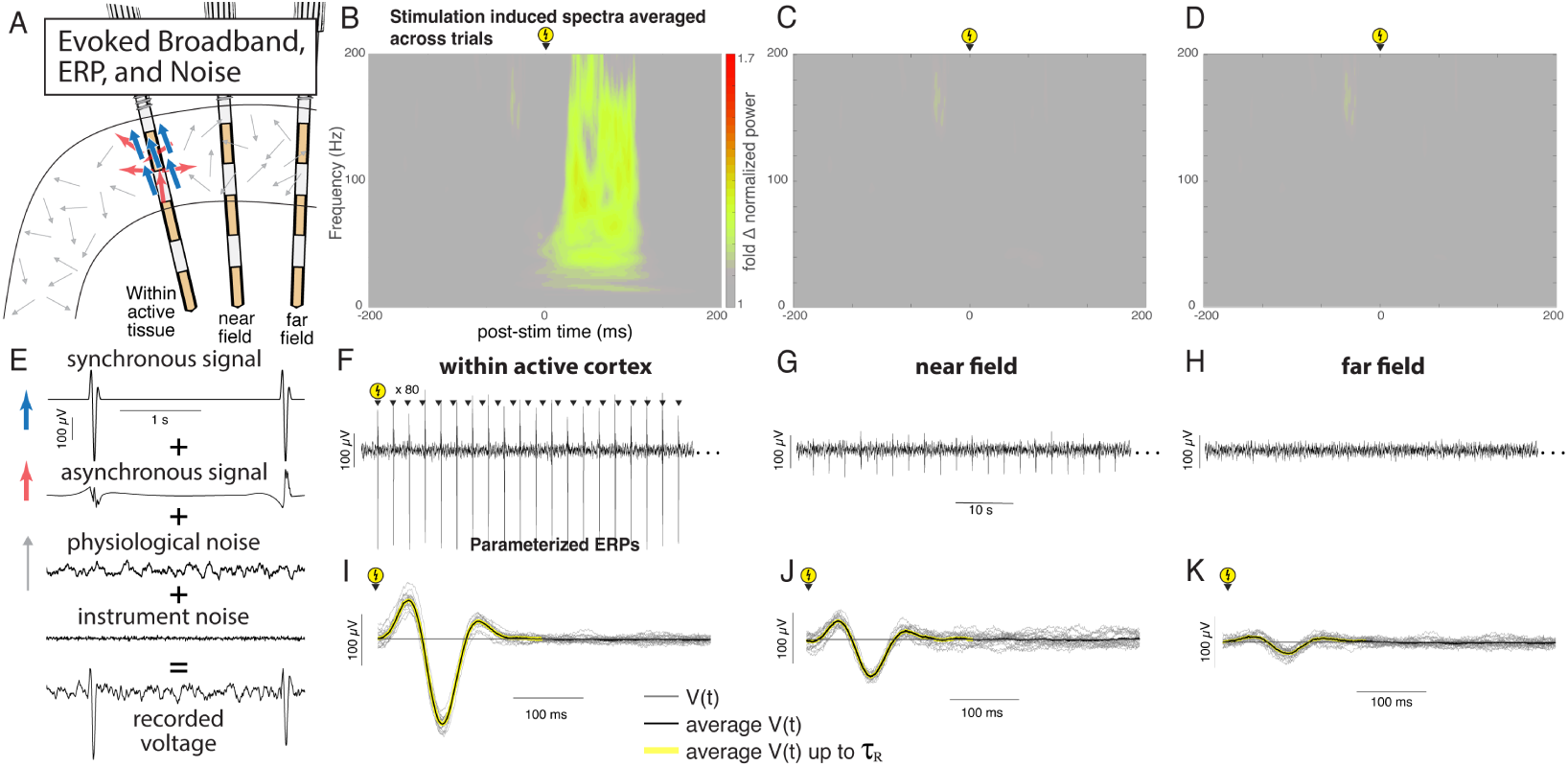
Synchronous signal (evoked potential) is more resolvable from background than asynchronous signal (evoked broadband). **A.** Schematic showing stimulation responsive tissue, with stimulation evoked increases in synchronous (blue) and asynchronous (red) firing within a background of noise (gray). **B.** Average stimulation induced spectrogram computed across 20 trials of the high SNR time series representing recording within active cortex. **C.** As in B, but computed from medium SNR time series representing a near field recording. **D.** As in B, but computed from low SNR time series representing a near field recording. **E.** Two example trials of the synchronous & asynchronous signals and the physiologic and instrument background noise combined to generate the synthetic data shown in F-H. **F-H.** Synthetic time series containing BSEPs and stim-evoked broadband within a background of noise. **I.** CRP parameterization of the time series representing recording within active tissue. **J.** As in I, but with a near field time series. **K.** As in I, but with far field time series. Note: a 200 ms significant response period is identified across all three noise levels while the stim-evoked broadband is progressively less resolvable

### Illustrating the effect of dipole source orientation on mCAR and bipolar signals

The orientation of a dipole to nearby recording channels will influence the sign and amplitude of mCAR and bipolar re-referenced BSEPs. In the brain, there are myriad orientations a dipole can take, but this principle is best understood using four cardinal orientations. 1) The dipole axis is perpendicular to and centered between two recordings electrodes (Fig 5a,b): the recorded voltage at each electrode will be identical, generating identical modified common average signals and a flat bipolar signal. This scenario is most often observed when the distance from the sources is much larger than the distance between adjacent electrodes. It is also possible when the source lies precisely perpendicular to the midpoint of two adjacent electrodes. 2) The dipole axis is perpendicular to and aligned to one recording electrode (Fig 5c,d): the recorded voltage of the mCAR and bipolar signals will have the same shape and sign but differ in magnitude. 3) The dipole axis is parallel and the equatorial plane is aligned to a single electrode (Fig 5e,f): the recorded voltage at the aligned electrode will be zero such that the bipolar signal is identical to the voltage recorded at the channel off of the equatorial plane. 4) The dipole axis is parallel and the equatorial plane centered between electrodes (Fig 5g,h): the mCAR signals are the same shape with opposite signs, leading to a bipolar signal with double the amplitude of the signal in each electrode.

**Fig 5.**
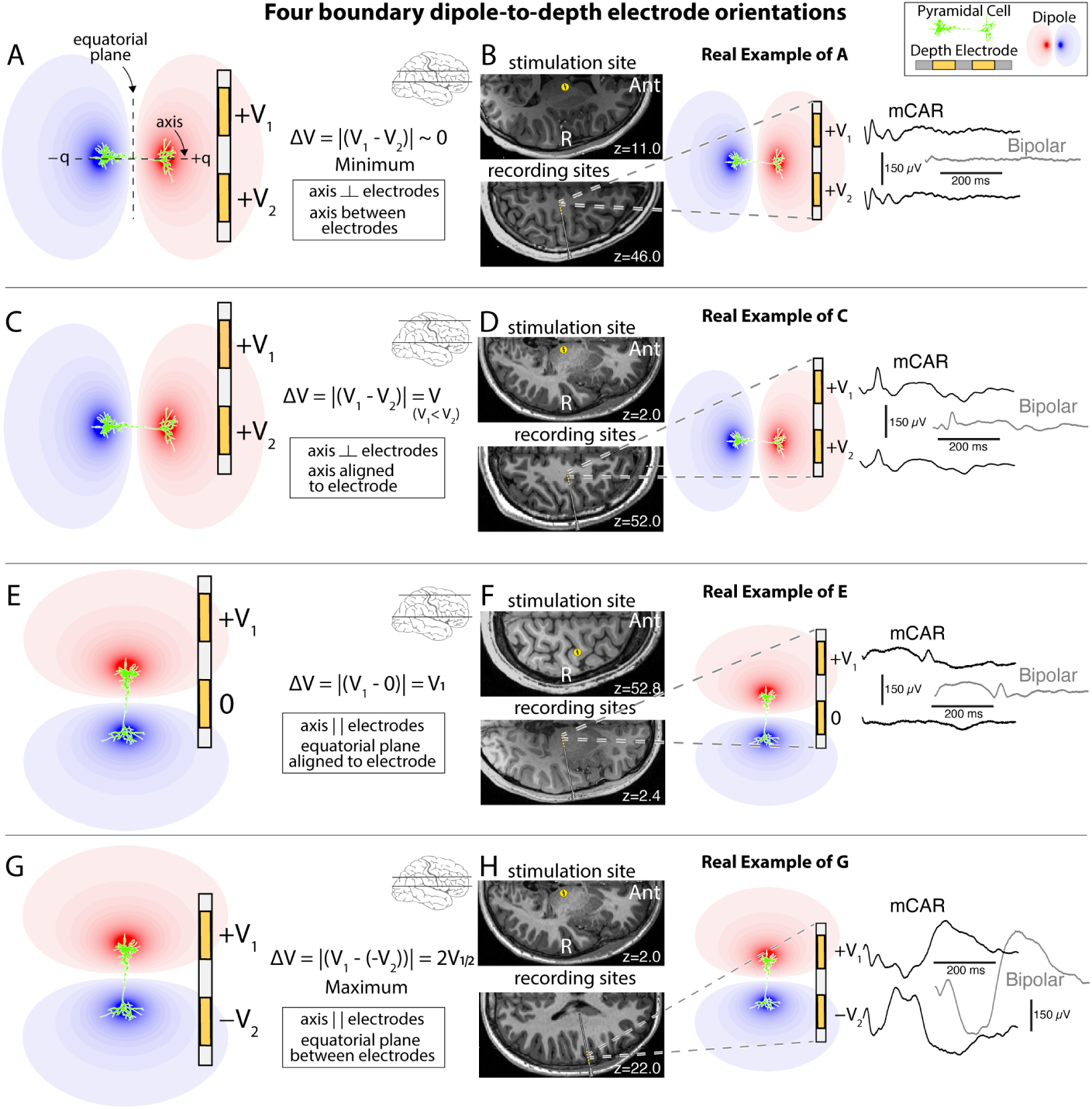
The orientation of bipolar channels to surrounding microcircuitry impacts voltage magnitude and may reflect dipole source orientation. **A.** When the dipole axis is perpendicular to the electrodes and sits halfway between electrodes, the difference between their recorded voltages is at a minimum (zero). **B.** An evoked potential from the right motor cortex is shown after stimulation in the right thalamus (data from subject 3). **C.** When the dipole axis is perpendicular to the electrodes and aligned to a single electrode, the difference between their recorded voltages is due to the difference in radial distances of each electrode to the dipole’s center. **D.** An evoked potential from the right motor cortex is shown after stimulation in the right thalamus (data from subject 2). **E.** When the dipole axis is parallel to the electrodes and the equatorial plane is aligned to a single electrode, the difference between their recorded voltages is equal to the voltage from the unaligned electrode as the fields cancel in the aligned electrode. **F.** An evoked potential from the right thalamus is shown after stimulation in the right motor cortex (data from subject 1). **G.** Schematic of a dipole generated across the long axis of a pyramidal cell. The axis of the dipole is coaxial to the pyramidal cell with the equatorial plane perpendicular, crossing the axis at its center. Axis is parallel to the electrodes and the equatorial plane sits halfway between electrodes, the difference between their recorded voltages is at a maximum. **H.** An evoked potential from the right temporal lobe is shown after stimulation in the right thalamus (data from subject 2).

### Estimation of the threshold distance to be able to record an evoked potential above ambient noise

At some distance, the recorded evoked potential falls beneath the aggregate noise. For example, in subject 3, cortical stimulation induced BSEPs of similar shape within two leads recording from the middle/anterior cingulate from a lateral approach, passing through white matter. As this shared shape likely represented a common source, the CRP method was applied to all mCAR signals in these two leads generating a single canonical shape with *α*’ values at each channel representing field strength (Fig 6b). These *α*’ values were then used to infer the location and orientation of the dipole source (see Methods for more details). Based on this inferred source position, the distance from the source at which the *α*’ values were no longer significant based on the CRP parameters were 18.3 and 20.2 mm for each lead, representing the approximate upper bound of the far field for this source (Fig 6c). Note that a 2 cm threshold is specific to this example, and does not represent a general boundary.

**Fig 6.**
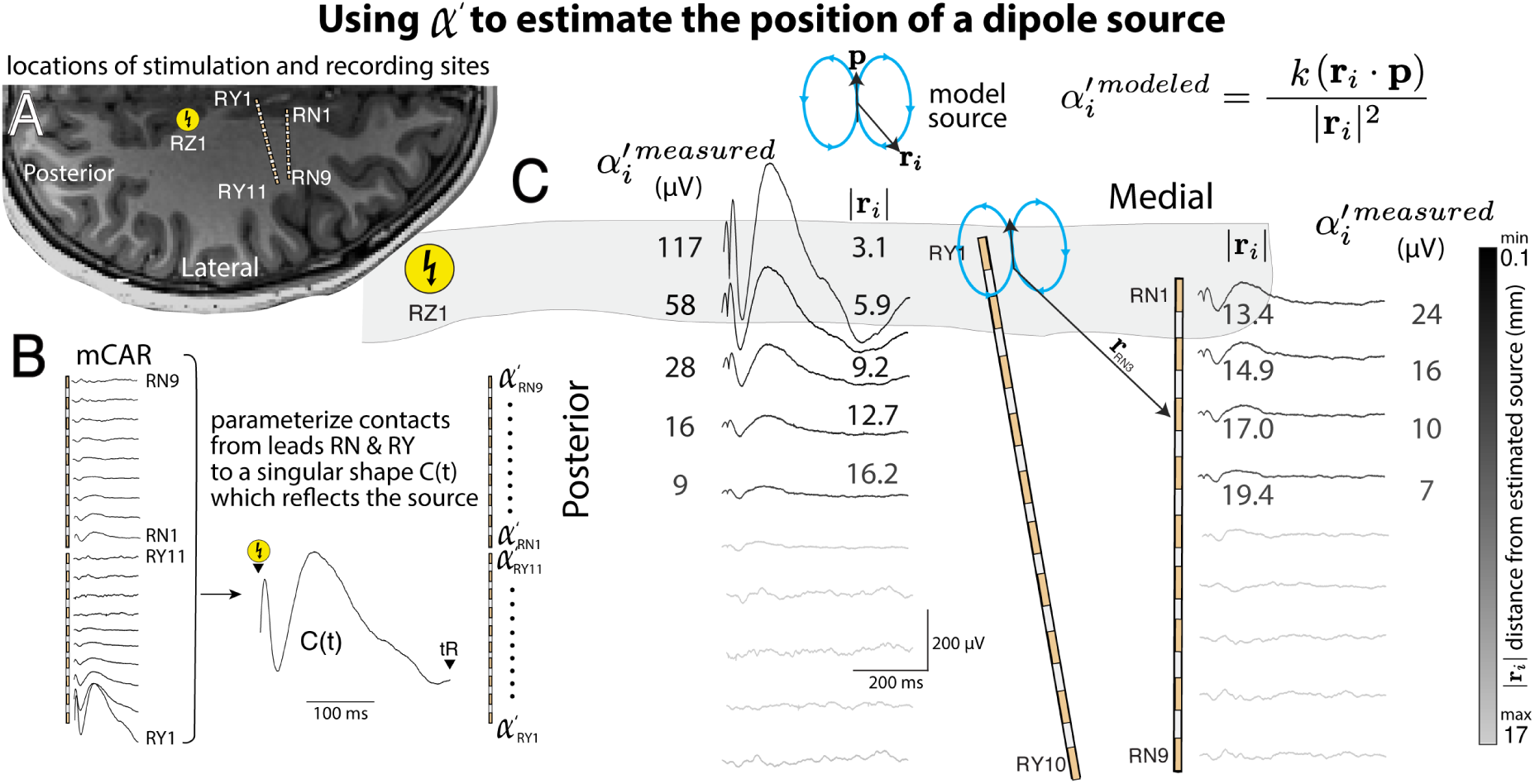
Illustration of a technique to empirically estimate the approximate position and magnitude of a dipole source. **A.** Two adjacent leads in the mid-cingulate recorded local field potentials near an induced source after stimulation at a site ≃ 2 cm away (RZ1). **B.** Recorded voltage time series across all trials from each nearby channel were parameterized together using the CRP method to generate a singular canonical response shape. The alpha prime (*α^′^*) values were used to estimate the strength of local field strength at each recording site. **C.** Schematic of the anatomical locations of recording sites in A, with associated *α^′^* values for each. All listed *α^′^* values are statistically significant (p *<* 0.05, one-sample t-test). The source position was estimated by minimizing the squared error between the measured voltages (mean *α^′^* at each recording site) and the modeled voltages. Gradient descent was used in find the minimum value until convergence criteria were met. Note that the actual inferred dipole projects approximately normal out of the page. Note that this is an inferred point source, but the actual source will have spatial complexity over a distributed region (Sup Fig 2)

### Mechanisms underlying BSEPs

#### Axonal mechanisms

The most prevelant hypothesis of how intracranial stimulation leads to a BSEP is via orthodromic propagation of action potentials along the white matter axons to other cortical region(s) monosynaptically (Fig 7a) or polysynaptically (Fig 7b) [20], but other plausible mechanisms exist. For example, stimulation may induce antidromic action potential transmission [21] (Fig 7c) or action potential propagation via intracortical axons (Fig 7d).

**Fig 7.**
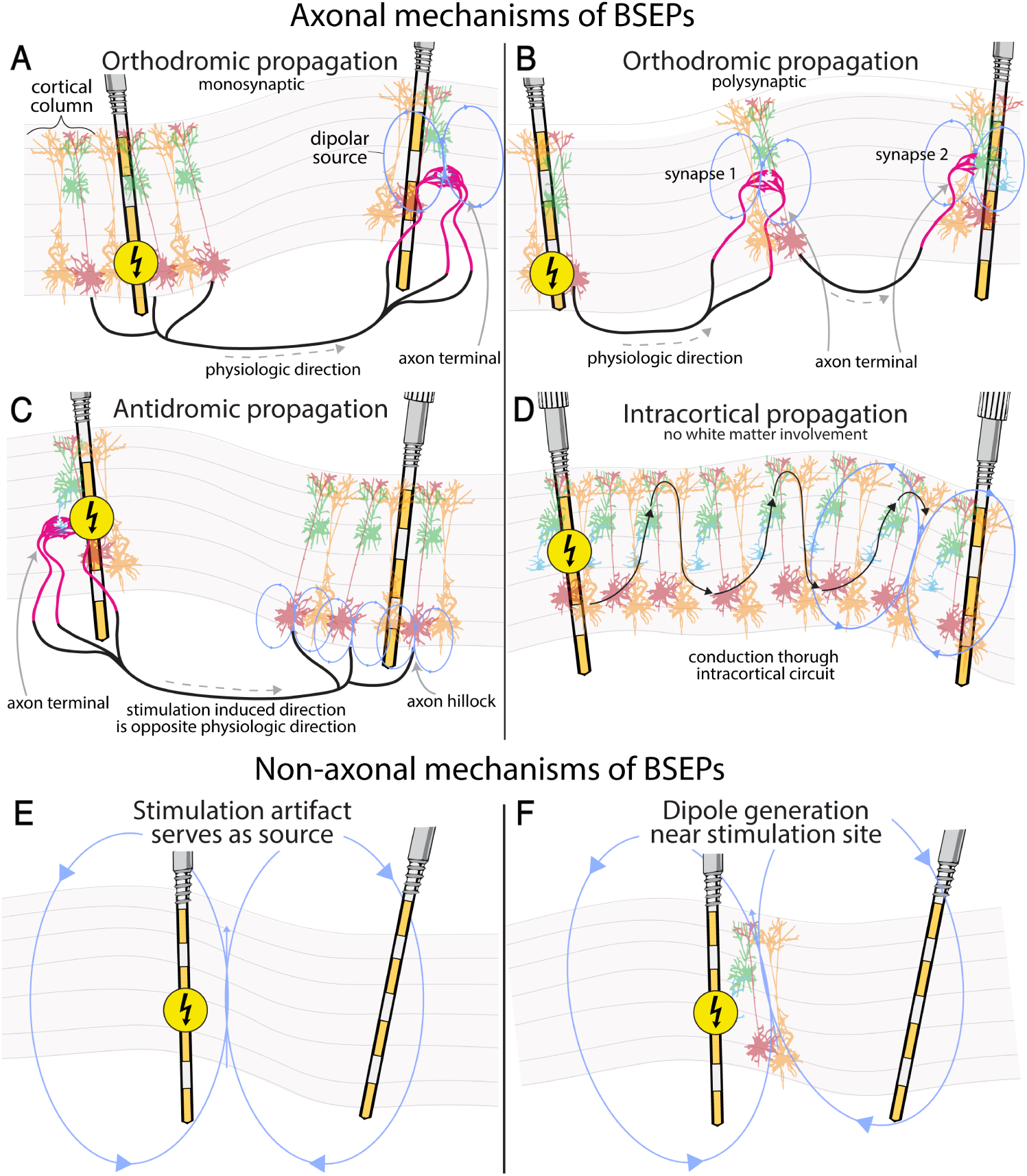
Axonal and non-axonal mechanisms of BSEPs. **A.** Stimulation may cause depolarization of surrounding tissue leading to orthodromic propagation of action potentials towards the circuitry surrounding a recording site (i.e. in a physiological direction). **B.** Stimulation induced orthodromic propagation of action potentials may also progress through multiple synpases pbefore arriving at the circuitry surrounding a recording site. **C.** Stimulation of axon terminals may lead to antidromic action potential propagation and depolarization of the upstream neurons. **D.** Stimulation in the cortex may lead to propagation of action potentials exclusively within the axons of gray matter towards the circuitry surrounding another cortical recording site. **Non-axonal mechanisms of BSEPs: recording near the stimulation site. E.** Stimulation – monopolar or bipolar - itself creates an electric field. **F.** Stimulation can induce a dipole in the tissue surrounding the stimulation site.

#### Non-axonal mechanisms

Bipolar stimulation will itself generate a field that can be measured. It can also depolarize adjacent tissue resulting in a neural source immediately around it (Fig 7e,f). As these sources are not due to axonal projections, they do not represent interactions between brain areas. Rejection of channels within a fixed radius of the stimulation site avoids false false interpretations related to this [7, 22].

### Stimulation-induced contamination of reference measurements

In intracranial EEG, white matter signals with low variation are often selected as a reference due to their relative stability compared to scalp references [23–25], but intracranial stimulation may challenge this stability. For example, stimulation that induces dipoles adjacent to the quiet white matter reference may contaminate the reference signal, introducing a measured global artifact in all channels (Fig 8 a,b). This pattern is seen as the same evoked potential shape in all recording channels (Fig 8c). Due to the consistency in the shape and amplitude of this artifact, bipolar or plain common average re-referencing cancels out this artifact in channels lacking a true response (Fig 8d).

**Fig 8.**
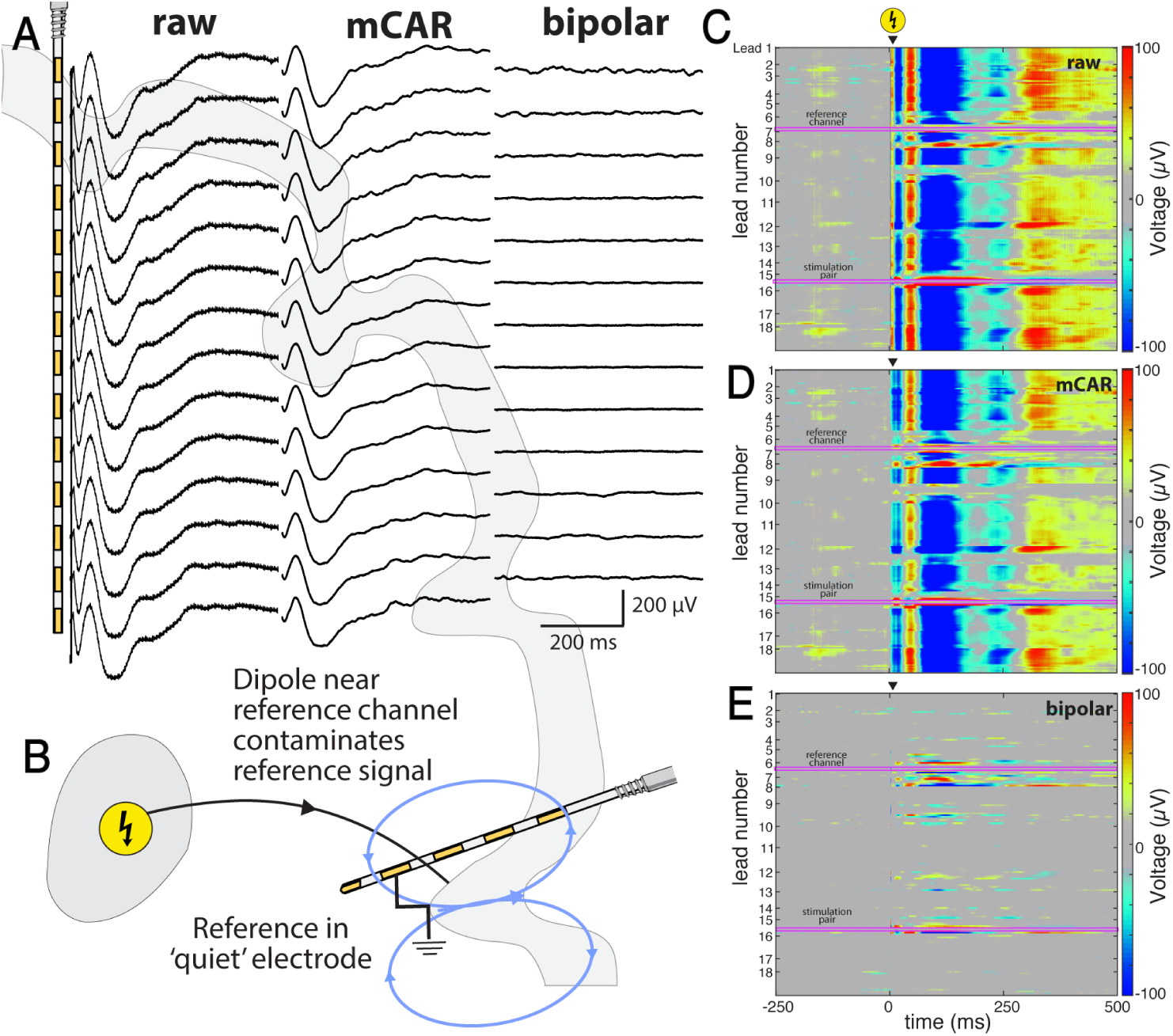
Dipoles near the reference channel create ubiquitous responses in all channels, (data from Subject 4). **A.** For certain stimulation pairs, one may measure a nearly identical voltage deflection in all channels if the reference channel is physically near an induced source. This will be observed in the raw (high-passed, notched) and modified common average re-referenced signal (mCAR), but no contamination is seen after bipolar re-referencing. **B.** A schematic illustrates how this pattern occurs when the intracranial reference channel sits near a stimulation induced dipole. **C.** Reference signal contamination is seen in the raw, non-re-referenced signal. **D.** mCAR re-referencing which uses low variance signal to construct a reference signal, fails to eliminate this contamination due to its high variance. **D.** Bipolar re-referencing eliminates the effect of contamination.

## Discussion

### Tailoring analysis to scientific questions based on 3 measurement domains

The aim of this study is to show how simple signal processing techniques can be used as tools to identify and characterize connections between stimulating and recording sites. Recording sites within active tissue contain both stimulation-evoked broadband power (BB) changes [26–28] alongside both bipolar and modified common average (mCAR) re-referenced BSEPs (Figs 2-4). As broadband measures local cortical activity [16], this result indicates that the stimulated site activates the circuitry at the recording site. This combination of BB and BSEPs is spatially precise and should be used when certainty regarding the relationship between stimulation and recording site is desired as when localizing potential targets of neuromodulation such as deep brain stimulation.

In the absence of BB change, BSEPs alone can be used to quantify connectivity [2, 5, 26]. When recording near field, BB changes are absent while bipolar and modified common average (mCAR) re-referenced BSEPs are present (Figs 2-4). In these cases, re-referencing can be leveraged as a tool. When the assessment and quantification of connectivity is the sole aim of a study, bipolar re-referencing is useful as it limits identification of channels to the near field, still allowing for the measurement of connectivity strength. As bipolar re-referenced BSEPs can vary in shape due to the orientation of the dipoles they measure (Fig 5), mCAR re-referencing is preferred when examining BSEP morphology (e.g. early voltage deflections). These near field recordings identify connectivity between the stimulation site and the region around a recording site, albeit with limited spatial resolution, and can be employed to define candidate regions for neuromodulation such as regional sensing applications.

Far field recordings lack BB and bipolar re-referenced BSEPs, but contain mCAR re-referenced BSEPs (Figs 2-4). This recording pattern establishes that the stimulation activates circuitry within the general region of a recording site. Although it does not map connectivity with high spatial resolution, it still allows for quantification of state changes (e.g. conduction speed) within the broader region of the recording site. These state changes can be used to assess the effects of interventions such as anesthesia or anti-seizure medication [29] (performing SPES before and after). Table 2 summarizes the potential interpretations and use cases for each measurement domain.

**Table 2.** Summary Characteristics for Each Measurement Domain. For each measurement domain, the response patterns (BB, BSEPs), valid interpretations, misinterpretations, use cases, and potential clinical applications are summarized. ^†^Note that there is a possibility that stimulation induces an effect with the same BB negative pattern but is near-field because it induces a synchronized response without activating circuitry. Besides antidromic action potential propagation, we have never seen an instance where we believed this to be the case. Evidence of this phenomenon would need to be identified with laminar recordings (e.g., the Neuropixel device [30]). *Baseline ERP required. ^ø^Note that direct includes both monosynaptic and polysynaptic connections. ‡Should be performed using mCAR signal to avoid confounders introduced by the dipole orientation’s influence on bipolar signal shape and magnitude. ^‡^Should be performed using mCAR signal to avoid confounders introduced by the dipole orientation’s influence on bipolar signal shape and magnitude. *Abbreviations:* BB = broadband power, BP = bipolar re-referenced BSEP, mCAR = modified common average re-referenced BSEP.

| Domain | BB | BP | mCAR | Potential interpretation | Potential misinterpretation | Optimal use case | Potential clinical use |
| --- | --- | --- | --- | --- | --- | --- | --- |
| Within Active Cortex | + | + | + | Stimulation activates circuitry at recording site | - | Establish direct influence<sup>o</sup> from stimulation to recording sites | Localize precise anatomical targets for neuromodulation (e.g., deep brain stimulation) |
| Near Field<sup>†</sup> | - | + | + | Stimulation activates circuitry nearby recording site (potential changes at smaller scale than inter-electrode spacing) | Stimulation activates circuitry at recording site. | Establish relationship between stimulation site and immediate region near recording sites<sup>o</sup>.<br>Quantify state changes<sup>‡</sup> (e.g., conduction speed, amplitude of BSEP) around recording site. | Define anatomical regions for neuromodulation (e.g., regional sensing applications) |
| Far Field | - | - | + | Stimulation activates circuitry more distant from recording site potential changes at larger scale than inter-electrode spacing) | Stimulation activates circuitry at recording site. Stimulation activates circuitry in immediate region of recording site. | Quantify state changes (e.g., conduction speed, amplitude of BSEP) in general region of recording site. | Assess overall state changes after intervention (e.g., chronic stimulation, tissue ablation, pharmacology). [29]<br>Effect of dynamically changing brain states (awake/asleep, cognitive task engagement, etc.) |

### Potential misinterpretations of BSEPs

While near field recordings provide valuable insights regarding the connectivity of the stimulation site to the region near a recording site, a lack of stimulation-evoked BB suggest that the source of the BSEP is not at the recording site. It is possible in some cases that a near field recording may sit within the tissue source but still lack BB, signifying that SPES-induced action potentials did not engage local circuitry, though we have never seen an instance where we believed this to be the case. Hypothetical instances of this response pattern might be: inhibitory inputs, antidromic inputs, inputs of insufficient magnitude to activate circuitry, BB responses too small to be recorded (from small volumes of gray matter), or scenarios when multiple inputs are required. Future studies involving laminar recordings (e.g. neuropixel device [30]) might be able to establish this mechanism. Far field recordings of mCAR BSEPs that don’t contain bipolar re-referenced BSEPs or stim-evoked BB do not demonstrate connectivity of the stimulation site with the recording site nor the nearby tissue as outlined above, and should not be misattributed to anatomical connectivity. See Table 2 for a summary of potential misinterpretations.

### Scientific constraints of near and far field recording sites

BSEPs may arise from stimulation-induced orthodromic or antidromic action potentials (Fig 7). This is because axonal depolarization travels both ways, and will potentially induce an effect whether current arrives at the axon terminals of a downstream synapse or axon hillock of an upstream neuron registering as a BSEP at nearby electrodes. Therefore, we cannot distinguish antidromic from orthodromic connections (Fig 7a-d). Similarly, the measurement domain of a recording site impacts how reciprocity between two sites [31, 32] can be interpreted. As near and far field recording sites do not sit within the circuitry generating the recorded BSEP, their stimulation may engage entirely different circuitry, confounding our interpretation of reciprocity. Sites within active cortex (BB+) avoid this constraint as the stimulated at and recorded from circuitry are the same.

### Consequences of reference contamination and solution

Intracranial white matter references can lack stability when contaminated by stimulation, even when stimulation is far from the reference channel but induces sources nearby (Fig 7b). Reference contamination must be identified by examining all responses for a single stimulation pair (Fig. 8c, d). Due to this, when BSEPs are studied in a single recording site in response to stimulation at several other sites (convergent analysis [26, 33]), reference contamination should be screened for as BSEPs assumed to be physiological may actually due to reference contamination. Prediction of which stimulation sites are more likely to impact white matter references would be useful (Supplementary Fig 4), but even without this knowledge, utilizing alternative intracranial white matter references is a simple solution [34]. As a rule, when bipolar re-referencing is not used, this artifact should be explicitly screened for.

### Quantification of domain boundaries

While we show the far field domain has an upper bound near 2 cm, this is just one example (Fig 6). A thorough quantification of the ranges of this boundary sorted by tissue type and stimulation parameters would require data across a larger cohort of channels (the same is true of the near field). Of note, in many (maybe the majority of) physiological cases, there are several distinct neural populations contributing to an evoked potential, such that a more nuanced model (e.g. mutipolar expansion) might be optimal. In practice, we suggest delineating the three domains on a case-by-case basis using these re-referencing approaches and spectral analysis tools.

## Conclusion

Recording sites of BSEPs can sit within, near, or far from the tissue origin of the potentials they measure. This distance impacts the questions that can be addressed regarding connectivity. We show that re-referencing and spectral analyses can be used as tools to understand the relationship between BSEPs and their sources.

## Supporting information

**Sup Fig 1. Stimulation and BSEP removal for spectral analysis.**

**Sup Fig 2. Four principle sources of artifact in stimulation evoked spectra.**

**Sup Fig 3. The impact of removing parameterized evoked response on spectra.**

**Sup Fig 4. The impact of a holding the parameterized shape length constant on spectra.**

**Sup Fig 5. Identification and removal of a contaminated reference.**

**Sup File. File containing Supplementary Methods, Results, and Discussion.**

## Acknowledgments

We wish to extend our gratitude to the patients who volunteered their time to participate in this research, to Bambi Wessel, Cindy Nelson, the staff at St. Mary’s hospital, and Peter Brunner for his critical assistance in the assembly of our electrophysiology rig and operations of the BCI2000 software. This work was supported by the German Ministry for Education and Research (BMBF) for BIFOLD (01IS18037A) - KRM the Institute of Information *\* Communications Technology Planning *\* Evaluation (IITP) grants funded by the Korea government (MSIT) (No. 2019-0-00079, Artificial Intelligence Graduate School Program, Korea University and No. 2022-0-00984, Development of Artificial Intelligence Technology for Personalized Plug-and-Play Explanation and Verification of Explanation) - KRM, NIH 1F31NS135898-01A - MAJ, NIH U01-NS128612 - KJM, Helene Houle Career Development Award. The contents of this manuscript are solely the responsibility of the authors and do not necessarily represent the official views of the NIH. Our funders did not play a role in study design, data collection and analysis, decision to publish, or preparation of the manuscript. “- three letter code” after each funding source refer to the associated authors.

