## supplementary material for "Approximating Signal Sources in Stereo-EEG Single Pulse Electrical Stimulation using Re-referencing and Spectral Analysis"

**DISCUSSION: Considerations in stimulation evoked spectra**

Stimulation evoked broadband power (BB) is a powerful tool when approximating the relative distance of recorded BSEPs to their tissue source. While several methods already exist to characterize stimulation-evoked voltage deflections, stimulation evoked BB power is less well studied, especially in the context of depth electrodes. This discussion is not meant to introduce a new method but to discuss the nuances of stimulation evoked spectral estimation.

*Minimizing artifacts & false positives*

When computing stimulation evoked BB, it is critical to avoid the introduction of false positives when computing power from the voltage times series. Upon developing the methods used in this article, four principle sources are seen across subjects and have been illustrated with example data in Supplementary Figure 2. *Artifact sources 1 and 2*: improper suppression of stimulation artifact and early (< 20 ms) sharp voltage deflections can approximate a discontinuity (i.e. edge effect) leading to artifact (Supplementary Figure 2 a,b). *Artifact sources 3 and 4*: Since BB represents uncorrelated activity, elimination of the correlated, synchronous activity represented in the evoked response of each trial can clean up the signal and optimize spectral estimation (discussed further below). We do this by parameterizing the evoked response for each trial in each recording channel using the CRP method and regressing this shape weighted to each of the “k” stimulation trials (⍺_k_C(t)) out of the raw voltage timeseries (V_k_(t)) after stimulation up until when the significance period ends. While this is meant to improve spectral estimation, it often confounds the results due to discontinuities introduced by the gap between the residual term (V_k_(t) - ⍺_k_C(t)) and the raw voltage at the points where parameterization begins and ends (Supplementary Figure 2c,d). By starting the parameterization at the point where signal is suppressed to zero (e.g. within the inverted Hann or “Bathtub” window) and by extending the end of parameterization such that the canonical shape equals zero at its end point, one can avoid this artifact and the false positives it can introduce.

*Regressing out the parameterized canonical shape*

One consideration is whether regression of a canonical shape is necessary. Supplementary Figure 3 shows that regression out of the shape impacts the number of significant results that are identified using a standard two sample t-test comparing power before and after stimulation. While this may not be the optimal test, it is critical to understand that a lack of regression of the evoked voltage response prior to spectral estimation can lead to a substantial change in the results due to the BSEP instead of the evoked BB from the residual term. This is still the case when spectral estimation is carried out above 60 Hz in an attempt to reduce the impact of the lower frequency components of the BSEP (Supplementary Figure 3 b,d).

*Effect of a fixed parametrization window on spectral estimation*

In certain cases, the time point at which the regressed out parameterized shape ends is a source of discontinuities between the raw voltage and the residual term. By adjusting the calculation of the parameterized significance period of our canonical shape, the gap between V_k_(t) and (V_k_(t) - ⍺_k_C(t)) can be reduced or eliminated. Unfortunately, for many channels, this gap cannot be avoided due to trial-by-trial variation of V_k_(t). One solution is to use a constant post-stimulation window when parameterizing the BSEP (Supplementary Figure 4a). The advantage of this decision is that artifact in the spectra can be sequestered to time points when evoked responses should have fell off, but its disadvantage is a less accurate parameterization of BSEPs. This lack of accuracy will lead to leakage of the BSEP into the residual term. *Due to this reduction in parameterization accuracy, we restrict its use to the context of spectral estimation and strongly urge scientist against this when parameterizing BSEPs*. Supplementary Figure 4 shows an example where fixing the parameterization period serves to avoid false positives due to discontinuities introduced by regression of the parameterized shape within our post-stimulation window of interest. Note that while fixing the parameterization period has little impact on the shape in this case (single trial of a single channel), this will certainly not the case across all trials and recording channels. Also note that a fixed parameterization window was *not* used in this manuscript nor is recommended, but is worth understanding to inform future methods.

**FIGURES**

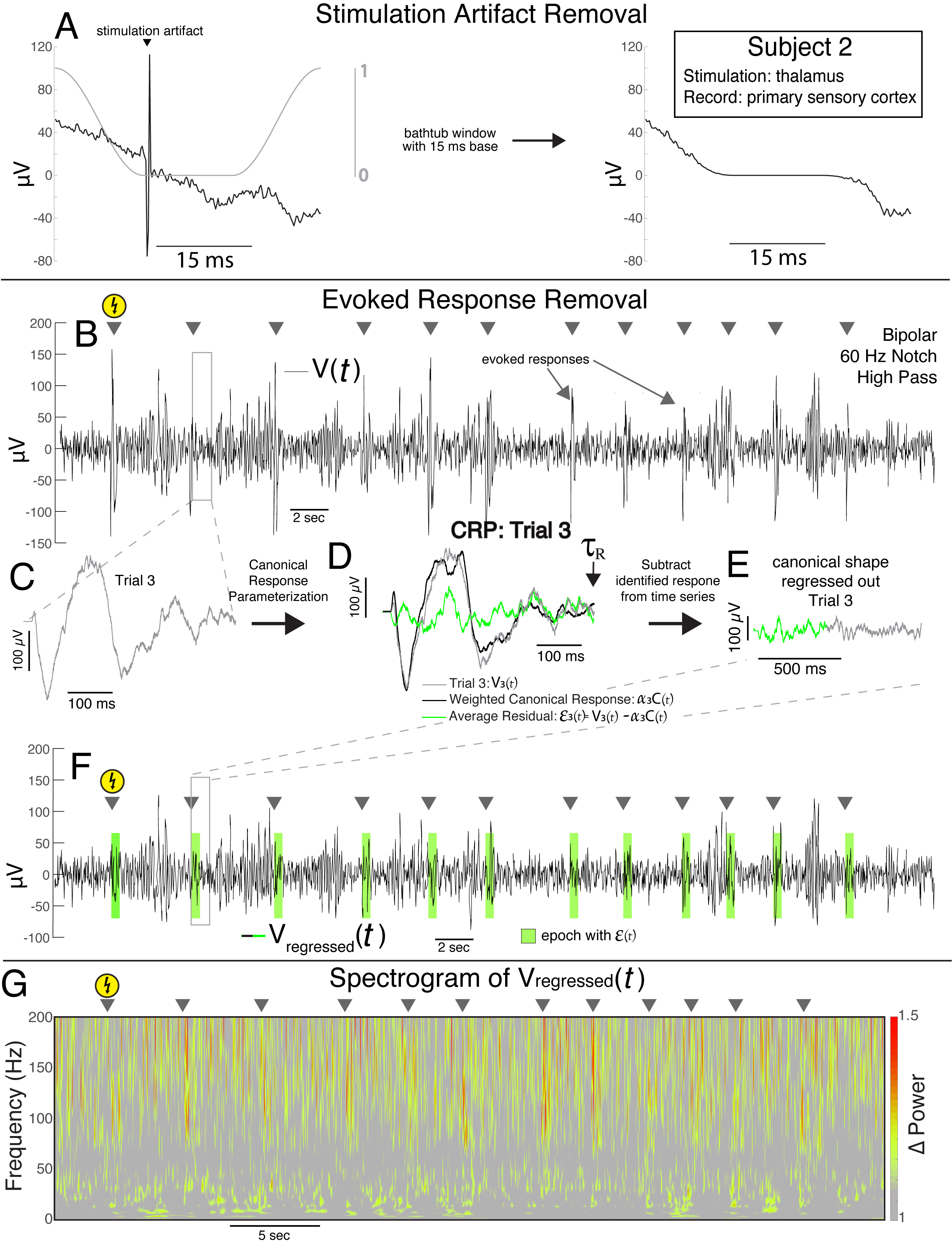

**Supplemental Figure 1. Stimulation and BSEP removal for spectral analysis. A.** In order to avoid the assumption that peri-stimulation signal contains is independent of stimulation, stimulation artifact was removed by multiplying an inverted Hann window to the 30 ms peri-stimulus period (15 ms before and after). **B.** Exemplary bipolar re-referenced data showing evoked response potentials (ERPs). **C.** ERP shown up to 1 s after stimulation trial 2. **C.** Recorded data following trial 2 before subtraction of the identified response. **D.** All data 15 ms to 2 s after stimulation were parameterized using the CRP method, which quantifies the significant response duration. **E.**  Recorded data following trial 2 after subtraction the identified response has been removed. Note only the significant time period is td. **F.** Entire experimental recording for a single stimulation pair after removal of identified responses. **G.** Spectrogram were computed by convolving a 5 cycle Morlet wavelet to the data from F. All data were normalized using the mean power for each frequency across the entire recording. Note this process identical when using CARLA method to re-reference after stimulation removal.

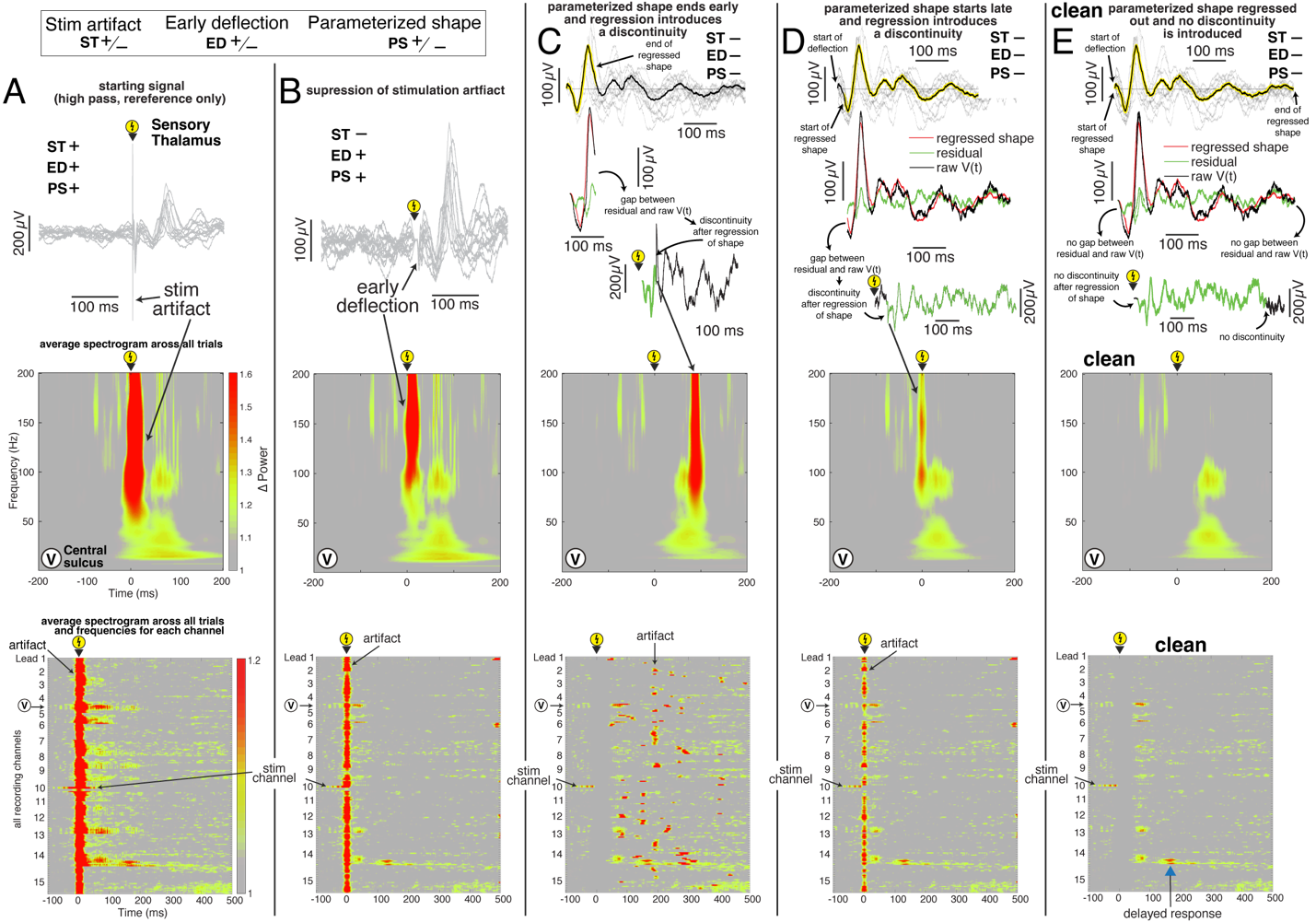

**Supplement Figure 2. Four principle sources of artifact in stimulation evoked spectra. A.** Stimulation artifact recorded from a channel in the central sulcus after stimulation in the sensory thalamus is evident as a sharp voltage deflection in all 12 voltage time series at the time of stimulation (top). This generates a large artifact at the time of stimulation in the evoked spectra averaged across trials centered at time zero (middle – recording in central sulcus). This pattern is present in all channels as seen in the evoked spectra averaged across trials and frequencies for each channel (bottom - arrows indicate the stimulation and recording channels in the top and middle panels above). **B.** Sharp deflections within the first 10 ms after stimulation also lead to artifact in the evoked spectra averaged across trials (middle) and may be a result of insufficient suppression of the stimulation artifact. This pattern is present in the majority of channels as seen in the evoked spectra averaged across trials and frequencies for each channel (bottom). Note: this artifact can be misattributed to stimulation. **C.** If the shape that is to be regressed out (e.g. parameterized shape or average shape) is ended prior to returning to baseline (~ 0 µV), a discontinuity is introduced due to the gap between the residual term, which is near 0 and the raw voltage prior to regression which is non-zero. This discontinuity leads to an artifact in the evoked spectra averaged across trials (middle) seen at the timepoint where the discontinuity is introduced. This pattern is present in all channels where the parameterized shape has been regressed out and can be seen in the evoked spectra averaged across trials and frequencies for each channel (bottom). **D.** As in C, a discontinuity can be introduced when the regressed shape that is to be regressed out starts after the initial voltage deflection such that there is a gap between the raw voltage and the residual term. This leads to an artifact in the evoked spectra averaged across trials (middle) at the timepoint where the discontinuity is introduced. This pattern is present in all channels where the parameterized shape has been regressed out and can be seen in the evoked spectra averaged across trials and frequencies for each channel (bottom). Note: this artifact can be misattributed to stimulation. **E.** By using the bathtub window to suppress stim artifact and early voltage deflections (Supp Figure 1a) and carefully avoiding discontinuities after regression of the parameterized shape, avoidable artifacts in the evoked spectra are removed. This is seen in the evoked spectra averaged across trials for a single channel recording in the central sulcus (middle) and evoked spectra averaged across trials and frequencies for each channel (bottom). Most evoked power can be seen between 50 and 100 ms with some exhibiting addition later responses (blue arrow).

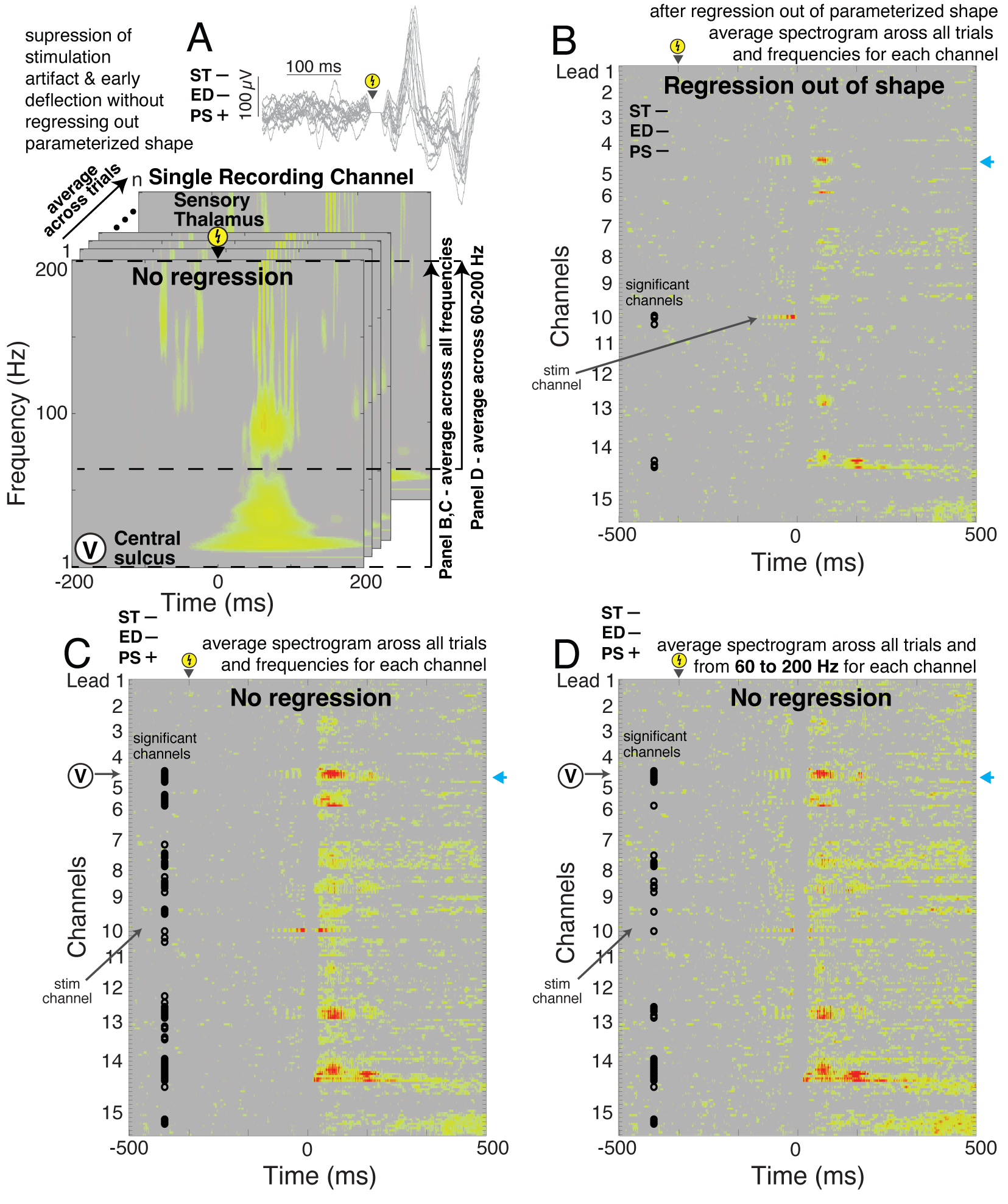

**Supplement Figure 3. The impact of removing parameterized evoked response on spectra. A.** The stimulation induced spectra (stimulation in sensory thalamus, recording in central sulcus) of an example channel - light blue arrows in B-D - is shown after stimulation artifact and early deflections have been removed using a 15 ms base bathtub window. In panel B & C, each row represents the average power across all frequencies (vertical arrow) and each trial (upper left corner arrow). In Panel D, each row represents the average power from 60 to 200 Hz (vertical arrow) and each trial (upper left corner arrow). **B.** Average spectrogram across trials and frequencies for all recording channels after stimulation of the sensory thalamus. Stim artifact, early sharp deflections, and the parameterized shape have been regressed out. Black circles indicate significant responses assessed by applying a two-sample t-test to the area under the curve of the power before (-500 to -20 ms) and after (20 to 500 ms) stimulation. P-values were corrected very strictly by multiplying by the number of recording channels (Bonferroni Method). **C.** As in B, but the parametrized shape is not regressed out. Note the increase in significant channels using the same significance criteria as in B. **D.** As in B, but the parametrized shape is not regressed out and power is only averaged from 60 to 200 Hz in an attempt to eliminate to limit the influence of the lower frequency components of the parameterized responses. Note the number of significant channels are increased from B, but fewer than C.

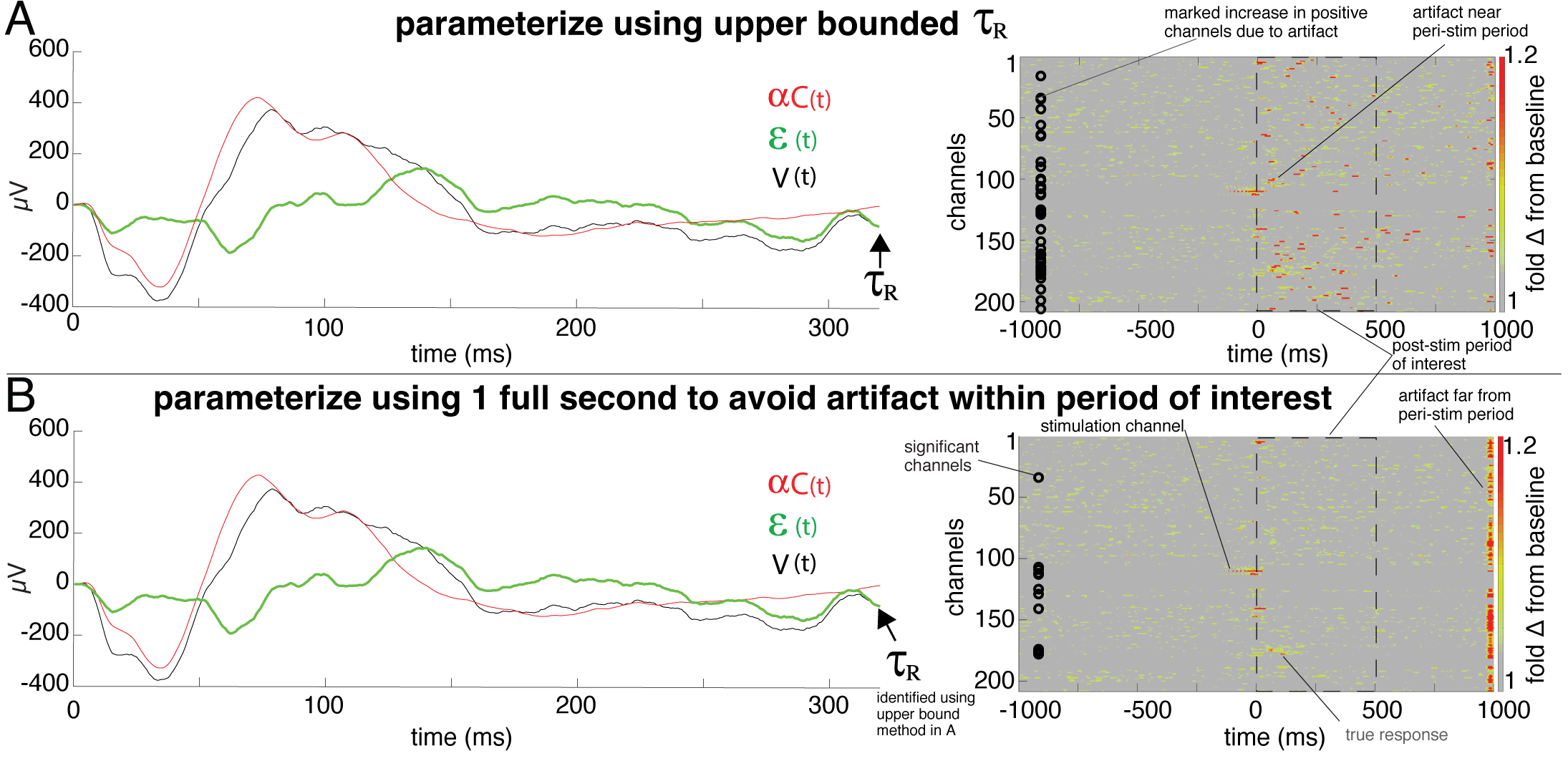

**Supplement Figure 4. The impact of a holding the parameterized shape length constant on spectra. A.** The raw voltage (black), weighted canonical shape (red), and residual (green) signals are shown in a recording site in the sensorimotor cortex for one stimulation trial in a site in the central sulcus (left). The significant period (tau R) was identified using the CRP method. CRP was applied to each channel and the parameterized shape was regressed out at the specific tau R identified for each channel. As seen in the traces (left), there is minimal difference between the parameterization whether tau R was used or not. In addition, the interference of artifact in the spectra as seen in B leads to many more false positive recording channels. **B.** As in A, but the period at which the voltage shape is parameterized was held constant at 1 second. As significance testing was performed only on data within 5 00 ms of stimulation, this avoided artifact within this window. Artifact is seen on the far right of the spectrogram where stitching in of the residual term leads to discontinuities. **Note**: even if parameterization differs slightly between scenarios A and B (using vs not using tau R), this less accurate parameterization will introduce some of the lower frequency component contained in the evoked voltage deflection. Influence of this on the spectrogram can be avoided by avoiding lower frequencies in the spectral analysis (e.g. using wavelets > 50 Hz). We do not recommend holding the parameterized shape constant, but think it helpful to the reader to illustrate.

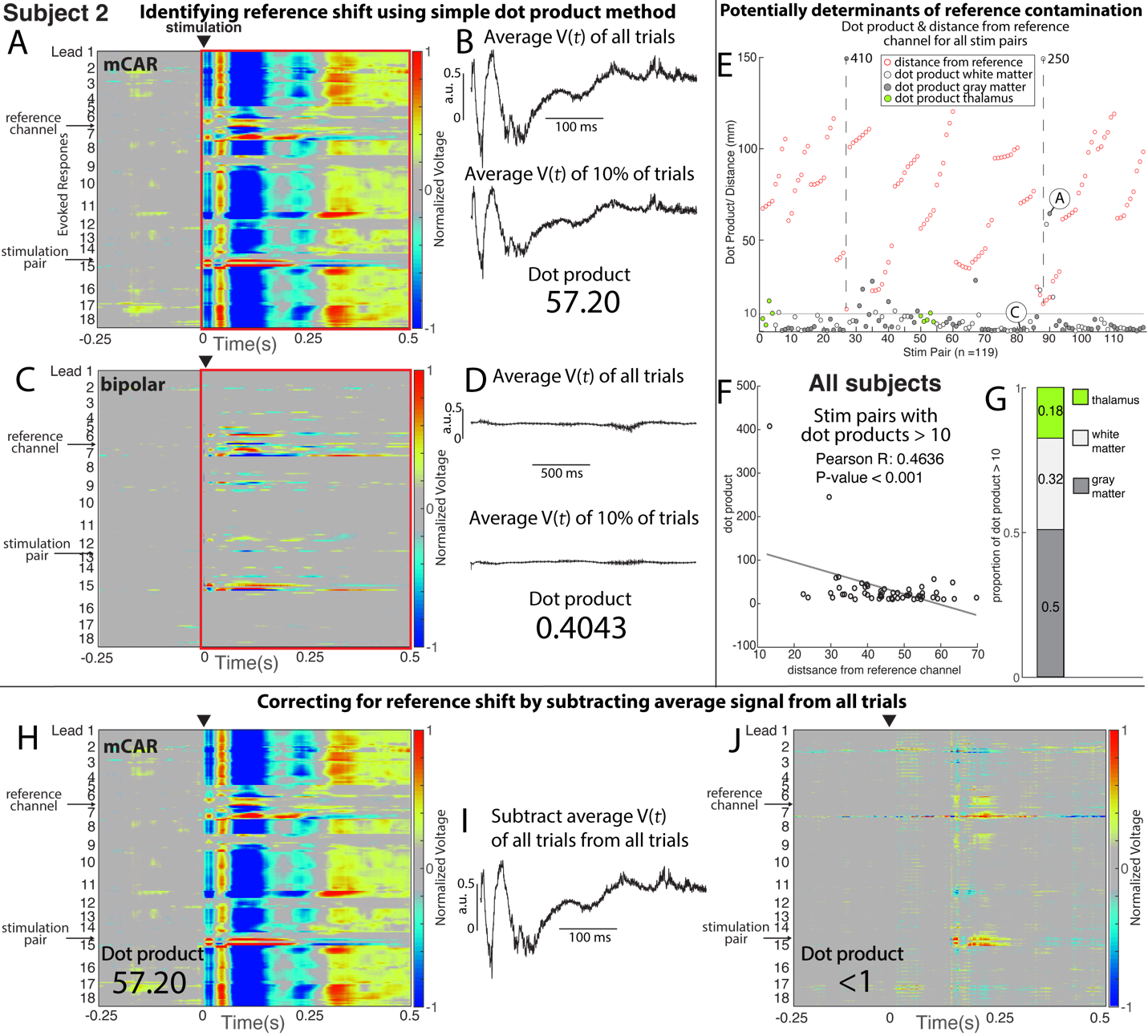

**Supplementary Figure 5. Identification and removal of a contaminated reference. A.** Evoked response heatmap showing all the response trials from 250 ms before to 500 ms after stimulation in the frontal lobe. Letter labels of the vertical axis (e.g. ‘RCM’) delineate the beginning and end of each recording lead. Arrows identify reference and stimulating channels. Note the strong responses in the channels surrounding the stimulation pair. **B.** The dot product between the average normalized response shape for the first 500 ms after stimulation (red box in A, C) across all trials in A (top) and a randomly selected subset of 10% of trials in A (bottom). **C.** As in A but during stimulation in the primary motor cortex. **D.** As in B, but with the responses shown in C. Note the much smaller magnitude of the normalized responses in D compared to B leading to a small dot product. **E.** The dot products for all stimulation pairs in Subject 3, with the fill color indicating the location of the interpolated position between each stimulated electrode pair: gray matter (dark gray), white matter (light gray), or the thalamus (green). Red circles indicate the Euclidean distance in millimeters between each stim pair and the reference channel. Note that the vertical axis indicates both the magnitude of the dot product metric and distance of the stimulation pair to the reference channel and the dot product metrics from A and B are indicated. **F.** For stimulation pairs with dot products above 10, there is a significance (p < 0.001, one sample t-test), albeit mild, positive relationship (correlation coefficient, r = 0.46) between the Euclidean distance of each stimulation to the reference channel and its dot product. **G.** Amongst all stimulation pairs with dot products above 10, 18%, 32%, and 50% are localized in the thalamus, gray matter, and white matter respectively. **H-J.** The evoked response heatmap from A (H) can be corrected by subtracting the average normalized response shape across all trials from each trial (I) to obtain clean responses (J).
